# Fast-growing *Bacillus sensu lato* rhizosphere populations are constrained by antagonistic Pseudomonadota, Actinomycetota *and other* Bacillus sensu lato

**DOI:** 10.1101/2025.09.17.676902

**Authors:** Muhammad Yasir Afzal, JaLeigha Kambeitz, Volker S Brozel

## Abstract

Copiotrophic *Bacillus* and related taxa grow rapidly and are commonly isolated from soil. Despite their growth rate, *Bacillus sensu lato* (BSL) constitute less than one percent of soil bacterial communities, and the nutrient-enriched rhizosphere contains even fewer. Amendment of bulk soil with synthetic root exudate did not lead to increase in *Bacillus* culturable counts. We hypothesized that BSL populations in soil enriched with growth-supporting carbon are suppressed by various soil microbes. A screen using *B. pseudomycoides* as tester strain yielded 124 growth inhibiting isolates, aligning by 16S rRNA genes to 3 *Alphaproteobacteria*, 6 *Betaproteobacteria*, 5 *Gammaproteobacteria*, 3 *Streptomyces*, and 19 *Bacillaceae*. Most antagonists also suppressed four other BSL, and over 70% of the BSL isolates suppressed each other. The 11 sequenced BSL genomes encoded between 2 and 10 antibiotic biosynthetic gene clusters. Incubation of multiple isolates in artificial soil microcosms resulted in population growth restraint through a high percentage of endospores formed. This indicated that growth suppression by antagonists was due primarily to induction of sporulation. These results support our hypothesis that *Bacillus* populations in soil enriched with growth-supporting carbon are suppressed by various soil microbes.

## Introduction

Fast-growing endospore-forming aerobic heterotrophs are among the best-known bacteria obtained from soils. These copiotrophs, widely called *Bacillus*, form large colonies on nutrient-rich agars within 24 h. Their ability to produce latent and extremely resilient cell structures called endospores enables them to survive in adverse environments [1–3]. Early microbiologists isolated *Bacillus* from soil regularly and considered them key members of the soil microbiota [4, 5]. Their omnipresence in soils has been confirmed through a meta-analysis of over 300 soil microbiome analyses [6]. However, they constitute only ∼0.4% of the soil microbiota. Species such as *B. subtilis* and *B. cereus* are known for their rapid growth, matched by high ribosomal operon copy numbers, 13 for *B. cereus* ATCC14579 [7]. Phylogenetic approaches have revealed a complex taxonomy for the classical *Bacillus* genus, which have over time been spread across several families of *Bacillota* (formerly *Firmicutes*). The aerobic Gram–positive endospore-formers have been assigned to *Alicyclobacillaceae*, *Bacillaceae*, *Cytobacillaceae*, *Caryophanaceae*, and *Paenibacillaceae* of the order *Caryophanales* [8, 9]. Yet the *Caryophanales* also contain non-spore-forming bacteria such as *Listeria* and *Staphylococcus*, so that there is no formal taxonomic name that incorporates all aerobic Gram–positive endospore-formers. We propose the term *Bacillus sensu lato* (BSL) for the spore-forming *Caryophanales*. The term *sensu lato*, meaning “in the broader sense” is already well established for the group of species closely related to *B. cereus*, widely termed “*B. cereus sensu lato*” [10]. The order *Caryophanales*, predominated by BSL, contain on average, eight ribosomal operon copies (obtained from rrnDB at https://rrndb.umms.med.umich.edu/, last accessed on 07/10/2025) [11]. These data predict rapid growth rates for members of the BSL.

A key feature of BSL is their ability to metabolize a wide range of carbon (C) compounds that support rapid growth [12]. C availability is often a limiting factor for microbial growth in soil, especially for copiotrophic BSL, which readily form endospores in response to nutrient limitation or other signals [13–15]. Several C sources act as environmental cues, leading spores to germinate into vegetative cells. This ability to sporulate and germinate later in response to specific cues is crucial for the ecological success of BSL [16, 17]. While soils are generally limited in terms of labile organic carbon, the region near roots contains C-rich exudates and rhizodeposits [18], and fast-growing taxa increase in abundance, which is termed the “rhizosphere effect” [19]. Growth rate has been proposed as an indicator of bacterial success in the carbon-enriched rhizosphere [20]. Recent tools to estimate the growth rate or Predicted Minimal Doubling Time (PMDT) from genome sequences [21] have enabled the testing of this hypothesis. A comparison of metagenomes from paired rhizosphere and bulk soil datasets supported the hypothesis that growth rate potential contributes to the rhizosphere effect by enriching genera with shorter PMDTs [22]. The exceptions to this pattern were *Firmicutes* (*Bacillota*). Both the bulk soil and rhizosphere MAGs (metagenome-assembled genomes) of *Firmicutes* predicted very short PMDTs, in contrast to all other phyla/classes, confirming that these species are fast-growing (see Fig. 3 in Lopez *et al*.) [22]. However, *Firmicutes* were the one phylum not enriched in the rhizosphere. These findings are supported by multiple microbiome analyses where BSL populations did not always show appreciable increases in the nutrient-rich rhizosphere in unsterilized soils [23]. A meta-analysis of paired clone libraries of bulk soil versus rhizosphere indicated a lower abundance of *Firmicutes* in the rhizosphere [20]. *Firmicutes* decreased from 10% in bulk soil to 5% in the soybean rhizosphere, indicating decreased fitness compared with that of *Proteobacteria* and *Bacteroidetes* [24]. Soil amendment with manure led to an increase in *Proteobacteria* but a decrease in *Firmicutes* [25]. We also reported that *Bacillus cereus sensu lato* populations did not increase when bulk soil was supplemented with a synthetic root exudate cocktail [26]. Collectively, these data suggest that BSL in the rhizosphere experience population-limiting pressure in the presence of labile organic carbon.

Soil bacterial communities are highly dynamic, with bacterial populations interacting with each other through cooperation, competition, and chemical signaling. Many factors can hinder the growth and proliferation of BSL in soils, including competition with other microbes for nutrition, biotic and abiotic stresses, bacterial viruses, and sporulation [27–29]. The soil and rhizosphere contain diverse microbial communities that compete for space, nutrients, and other resources. Some are involved in chemical warfare. Several gram-positive and gram-negative bacteria, such as *Streptomyces* (*Actinomycetes*) and *Pseudomonas*, are rivals of *Bacillus*, and certain fungi also produce compounds that are bacteriostatic or bactericidal to other bacteria, including *Bacillus* [30–33]. For example, *Streptomyces aureofasciens* produces tetracycline [34], *Penicillium chrysogenum* produces penicillin [35, 36], and *Pseudomonas aeruginosa* and *Pseudomonas fluorescens* produce hydrogen cyanide [37] which hinder the growth of other bacteria, including *Bacillus*. Toxic compounds produced by *P. fluorescens* are linked to nutrient composition, which leads to the disappearance of *B. thuringiensis* [38, 39]. These findings indicate that other soil bacteria likely limit BSL populations in the rhizosphere through production of suppressing compounds.

Despite the abundance of labile organic carbon in rhizosphere [40] and nutrient-amended soils [26], the number of *Bacillus* did not increase as expected, leading us to investigate this phenomenon. We hypothesized that BSL populations in soil enriched with growth-supporting carbon are suppressed by various soil microbes. While viruses and other microorganisms can also affect BSL populations, we focus here on fast-growing bacteria as potential antagonists. Our study revealed antagonism by diverse *Pseudomonadota* and *Streptomyces* but also by other BSL. Surprisingly, many of the BSL isolates inhibited each other. These results reveal the complex competitive relationships within the BSL group, which are influenced by conflicts within their own group.

## Materials and methods

### Determining the prevalence of BSL in soybean monocultures

To obtain culture-independent data on the population sizes of *Caryophanales* and specific BSL in bulk soil versus the rhizosphere under monoculture conditions of annual crops, we extracted data from our recent analysis of the bacteriota in the soybean rhizosphere and surrounding bulk soil from three sites in South Dakota [41]. Each site was sampled and amplified four times, and the amplified V1--9 region was sequenced by Oxford Nanopore Technology (ONT). We extracted the proportions of the bacterial communities of *Bacillota, Bacilli,* and *Caryophanales* and of the ten most abundant species of BSL from the abundance tables and plotted the data via GraphPad Prism 10 (Boston, MA, USA).

### BSL population prevalence in nutrient-amended soils

To assess the response of fast-growing total aerobic heterotrophs to root exudates, *B. cereus sensu lato* were enumerated in soils under different cropping conditions. Samples were collected from soybean (44°17’47″ N, 96°39’24″ W), corn (44°18’33.7″ N, 96°40’28.2″ W), and wheat fields (44°18’30″ N, 96°40’02″ W) and from a natural grassland (44° 2’10″ N, 96°47’6″ W). The samples were taken to a depth of 14 cm and placed in sterile bags for immediate transportation to the laboratory. Twenty grams of soil was homogenized with 2 mL of synthetic root exudate cocktail (REC) [26] using a sterile fork at time 0 and again after 24 h in the same soil. The culturable counts were determined at 0, 12, 24, 36, and 48 h after supplementation with REC at time 0 and after 24 h. Suspensions were prepared by adding 1 g of soil to 9 mL of high-purity sterile water and homogenizing by vortexing three times for 3 s. Serial dilutions (100 μL) were spread on triplicate plates of R2A (1.5% agar) (Research Products International Cat R24200, Mt Prospect, USA) and Mannitol Egg Yolk Polymyxin (MYP) agar (1.5% agar) (Oxoid catalog CM0929, Basingstoke, UK) supplemented with 10,000 units of polymyxin per L (Oxoid catalog SR0099E, Basingstoke, UK) and sterile egg yolk emulsion. Endospores were enumerated by treating the same dilutions at 80 °C for 10 min to kill the vegetative cells and plating them on R2A and MYP agars. The agar plates were incubated for 24 h at 30 °C, after which the colonies were counted.

To determine the population dynamics of *B. cereus* in the absence of other microbiota, we inoculated *B. cereus* ATCC 14579 into sterilized soybean soil and artificial soil. The soybean field soil (20 g) was autoclaved for two liquid cycles and again for one dry cycle to ensure sterility. The soil was supplemented with 500 µl of *B. cereus* ATCC 14579, grown overnight in R2A broth and incubated at room temperature (21 °C). After 36 h, a further 250 µl of fresh overnight-grown *B. cereus* culture was added, together with 2 mL of REC, t0, for this experiment. The dilutions were plated onto R2A at t0 and 12, 24, 36, and 48 h and incubated at 30 °C for 24 h.

To evaluate *B. cereus* in soil with no prior microbial history, artificial soil microcosms (ASMs) were prepared as described previously [42, 43], but supplemented with REC in place of the soil extract or LB and autoclaved for two dry cycles. The soil microcosms were inoculated and supplemented with REC as described above for sterilized soil. The culturable counts were determined as for sterilized soil.

### Enumeration of culturable antagonists

To determine the prevalence of rapidly growing antagonists, serially diluted suspensions of the four soils (10 μL) were spotted adjacent to spots of *B. pseudomycoides* on square plates containing R2A (9 spots of *B. pseudomycoides* and 16 spots of soil suspensions). We chose *B. pseudomycoides* because it quickly expands across the agar surface, facilitating easy visualization of growth suppression. This isolate was obtained from one of the soil samples, and its phylogeny was determined by sequencing the 16S rRNA gene. Local suppression of *B. pseudomycoides* growth indicated the presence of at least one antagonistic strain in the diluted soil aliquot.

### Isolation of potential antagonists of B. cereus sensu lato

Serially diluted suspensions of the four soil types were plated on R2A (1.5% agar) and incubated for 24 h at 30 °C. The plates were then overlaid with soft R2A (0.75% agar) containing exponentially growing *B. pseudomycoides* and incubated for another 24 h at 30 °C. Clear zones indicated potential antagonist colonies on the underlying agar. Isolation of the underlying colony required suppression of the rapidly growing *B*. *pseudomycoides*. To suppress *B. pseudomycoides,* we used crystal violet, which is known to inhibit the growth of gram-positive bacteria. The concentration of crystal violet that suppressed *B. pseudomycoides* without repressing the growth of other gram-positive bacteria was found to be 10 times lower than that employed in MacConkey and Eosin Methylene Blue (EMB) agars [44, 45] or 0.2 µg/mL. Potential antagonists were confirmed by streaking a line across agar and a line of *B*. *pseudomycoides* in a V-shaped fashion, and of these, 124 isolates were confirmed as distance-acting antagonists. The suppression phenomenon was also determined against four other BSL*, B. subtilis* 168*, Paenibacillus polymyxa* ATCC 48*, Priestia megaterium* ATCC 14581 *and B. cereus* ATCC 14579, via the cross-streak method.

### Determining the phylogeny of isolates by 16s rRNA gene sequencing

DNA was extracted via the Quick DNA ^TM^ MiniPrep Kit (Zymo Cat. No. D3024, USA), and the 16S rRNA gene was amplified via the universal primers 27F: 5’-AGAGTTTGATCCTGGCTCAG-3’ and 1492R: 5’-TACGGYTACCTTGTTACGACTT-3’ [46]. The PCR amplicons were submitted for full-length sequencing by Plasmidsauris (Arcadia, CA, USA). The 16S rRNA genes of related bacteria were obtained via the highly similar sequences (megablast) option in BLAST [47] and aligned to isolate sequences via MUSCLE [48]. The phylogeny was inferred using the Maximum Likelihood method and Tamura-Nei model of nucleotide substitutions [49] and the tree with the highest log likelihood (-8,557.09) is shown. The percentage of replicate trees in which the associated taxa clustered together (100 replicates) is shown next to the branches [50]. The initial tree for the heuristic search was selected by choosing the tree with the superior log-likelihood between a Neighbor-Joining (NJ) tree [51] and a Maximum Parsimony (MP) tree. The NJ tree was generated using a matrix of pairwise distances computed using the Tamura-Nei (1993) model [1]. The MP tree had the shortest length among 10 MP tree searches, each performed with a randomly generated starting tree. The analytical procedure encompassed 71 nucleotide sequences with 1,478 positions in the final dataset. Evolutionary analyses were conducted in MEGA12 [52] utilizing up to 4 parallel computing threads.

The gram-positive and gram-negative isolates were aligned separately. Because 16S rRNA gene sequences are not always able to differentiate among closely related species, we sought to determine species of the 11 sequenced genomes using three independent approaches: BLAST, Mash, and FastANI [47, 53, 54]. Mash compared the genome to large reference datasets to find the closest matches. FastANI then calculated precise genome-wide average nucleotide identity values, with 95 percent identity used as the accepted threshold for species identification. BLAST provided direct sequence alignments with high identity and coverage to a single reference species. Identification was based on agreement among the three approaches.

### Genome analysis of BSL isolates

The ten gram-positive isolates displaying the highest growth suppression activity and the *B. pseudomycoides* isolate were selected for whole-genome sequencing. DNA was extracted via two different methods. Isolates that were slimy in texture were extracted via the DNeasy UltraClean Microbial Kit (Qiagen Cat. No. D30241224-50, Hilden, Germany), but the bead beating was reduced to 5 instead of 10 min, and RNase (Thermo Fisher) was added after cell lysis. The remaining isolates were subjected to the chemical extraction method of “Wilson” [55] with some modifications to reduce DNA fragmentation. Zymo kit lysis buffer (Zymo Cat. No. D3024, USA) was added at step 1 to lyse the cells efficiently, and RNase was added at step 1 to degrade residual RNA. The DNA concentration and purity were determined via a Nanodrop spectrophotometer (Thermo Fisher Scientific, Pittsburgh, PA, USA). The genomes were sequenced via Oxford Nanopore Technology (ONT) at the SDSU Genomic Core Facility. The genomes were assembled via Flye. 2.9.4 [56], and BUSCO 5.8.2 [57] was used to determine the completeness of the genome assembly. Following assembly, the genomes were annotated via Prokka 1.14.6, and Prodigal 2.6.3 was used to predict the genes [58, 59]. After annotation, the genomes were submitted to the secondary metabolite biosynthesis gene cluster identification site antiSMASH 8.0 [60] to obtain evidence of antimicrobial biosynthesis-related gene clusters (BGCs).

### BSL population dynamics in the presence of putative antagonists in soil

To test whether isolates displaying antagonism on agar also suppressed the growth of BSL in soil and liquid microcosms, we co-inoculated one gram-negative isolate from each of the three classes (Fig. S1a, *Bosea robiniae, Pseudomonas chlororaphis*, and *Pseudoduganella guangdongensis*) together with *B. cereus* ATCC 14579 and separately four gram-positive isolates (*B. cereus*, *B. velezensis, P. megaterium,* and *P. simplex*). All the isolates were grown individually in R2B overnight, and equal numbers of cells (10^6^ CFU per ml) were suspended in 2 ml of REC and mixed with 20 g of sterile ASM. For the liquid microcosms, equal numbers of cells (10^6^ CFU per ml) were suspended in 5 ml of REC and 45 ml of sterile water. The microcosms were incubated at room temperature (21 °C), and serial dilutions were prepared at 0, 12, 24, 36, and 48 h. Dilutions of gram-negative and gram-positive bacteria before and after treatment at 80 °C for 10 min were plated onto MYP agar and R2A, respectively. *B. cereus* ATCC14579 inoculated on its own was enumerated via R2A.

## Results

### Prevalence of BSL in the bulk soil and rhizosphere

BSL appear to be less predominant in the rhizosphere than in bulk soil. To determine whether this holds true under monoculture conditions of annual crops, we extracted data from our recent analysis of the microbiota in the soybean rhizosphere and surrounding bulk soil [41]. The soil and rhizosphere used for DNA extraction were obtained from soybean plants at R3 or at the beginning of the pod stage. The phylum *Bacillota* was significantly less prevalent in the rhizosphere than in the bulk soil, as were the class *Bacilli* and the order *Caryophanales* (Fig. 1a). Eight of the ten most abundant species of BSL were also less predominant in the rhizosphere than in the bulk soil (Fig. 1b). These data show that BSL are less predominant in the nutrient-enriched rhizosphere of growing soybean plants and agree with several previous reports [24, 61, 62].

**Figure 1:**
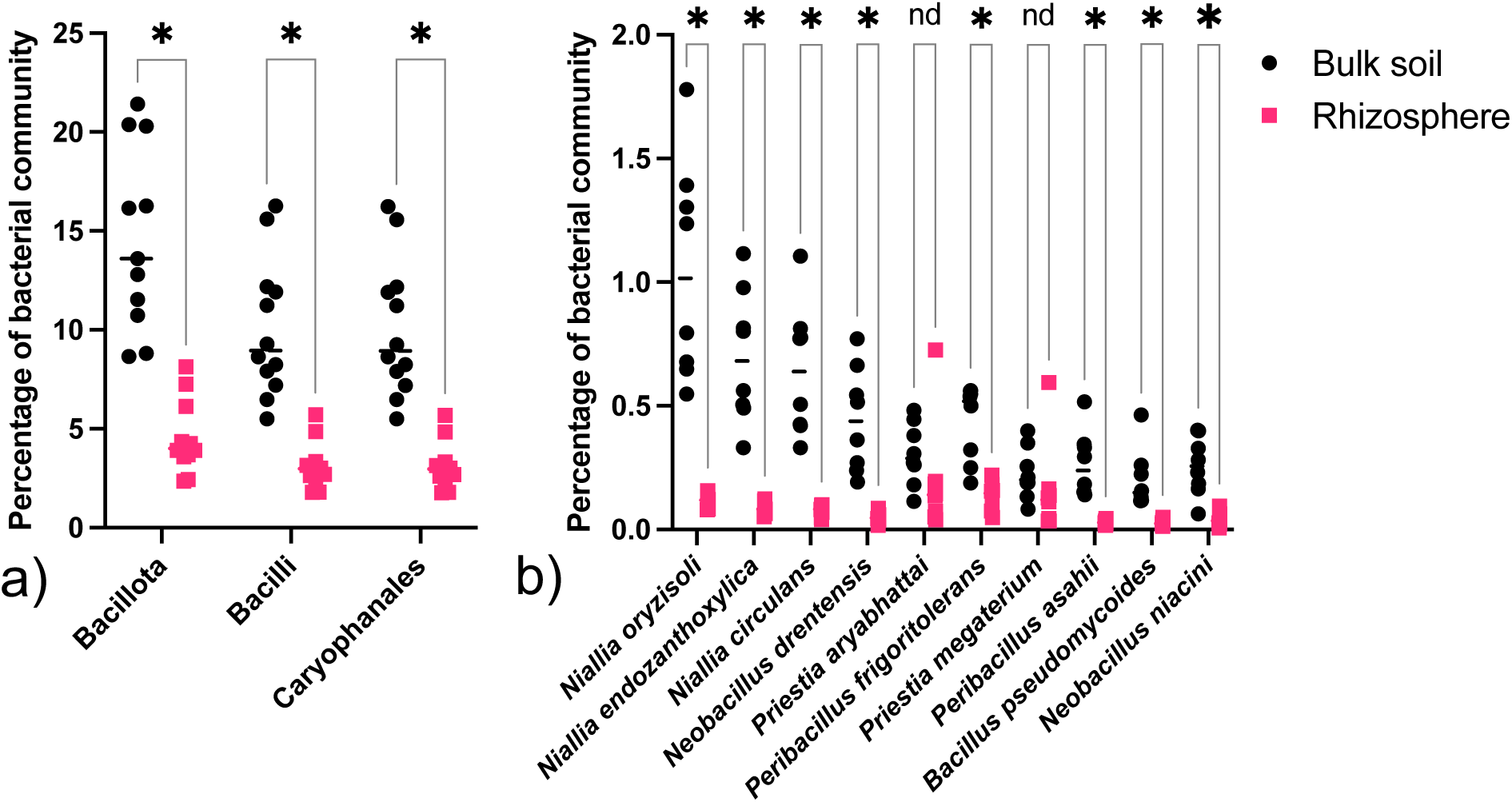
Proportions of the soil bacteriota in the bulk soil versus the soybean rhizosphere of members of the phylum *Bacillota*, class *Bacilli* and order *Caryophanales* (a), and the most abundant species of *Caryophanales* (b). Asterisks indicate significant differences (p<0.000001) between the bulk soil and rhizosphere samples via multiple unpaired t tests.

### BSL population prevalence in soils amended with exudate cocktail

To determine the population response of BSL in soil to the availability of root exudates, we determined the culturable count of *B. cereus sensu lato,* as this is the only group of BSL for which a selective medium is available [63, 64]. The soil samples were amended with REC, and *B. cereus* vegetative cells and endospores were enumerated via MYP agar. BSL, such as *B. cereus* and *B. pseudomycoides,* grow rapidly on agar when nutrients become available (Fig. 2a). The overall aerobic culturable count in the soil increased 100-fold following REC introduction, but the numbers of *B. cereus* remained unchanged (Fig. 2 b, c, d, e). This lack of increase in *B. cereus* was observed for all four soil types studied. Interestingly, the number of endospores also did not change, indicating either a lack of germination or germination followed by sporulation within 12 h. Modifications to REC, such as the exclusion of individual or all organic acids, yielded the same lack of change in the number of *B. cereus* in the soils (data not shown).

**Figure 2.**
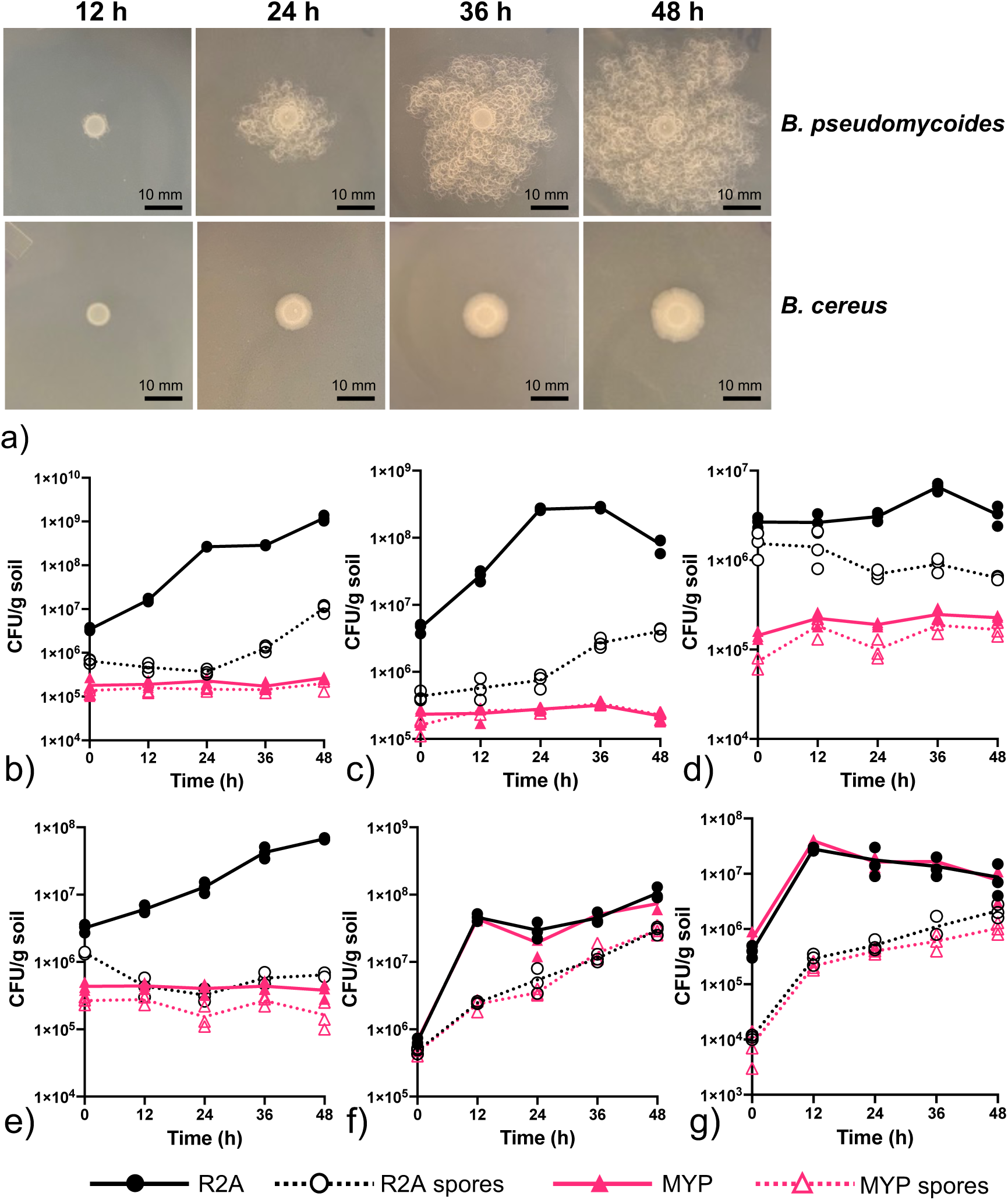
Colony development of *B. pseudomycoides* and *B. cereus* on R2A agar at 30 °C (a). Overall culturable aerobic heterotrophs and spores on R2A agar and *B. cereus sensu lato* total and spore counts on MYP agar in soybean field (b), wheat field (c), corn field (d) and grassland (e) soils and *B. cereus* inoculated into sterilized soybean field soil (f) and into synthetic soil with no prior history of bacterial presence (g).

To determine whether the resident microbiota was associated with a lack of BSL growth, we inoculated *B. cereus* into sterilized soybean fields and virgin soil (ASM) 36 h prior to the start of the experiment. After 36 h of population establishment (t0), the total and endospore counts indicated that all the cells inoculated into the sterilized soil had sporulated (Fig. 2f, t0), whereas the cells in the virgin soil remained mostly vegetative (Fig. 2g, t0). These findings suggest that compounds associated with microbial activity in soil prior to autoclaving induce sporulation. Importantly, both populations increased over 10-fold within 12 h of exposure to REC. The absence of other microbiota in the soil was associated with an increase in the population of *B. cereus*. These results indicate that the presence of other microbial communities in natural soils is associated with the inhibition of BSL and with the suppression of germination.

### Quantification of bacteria antagonistic to BSL

To determine the number of rapidly growing soil bacteria that suppress the growth of BSL, we spotted diluted soil suspensions near spots of rapidly expanding *B. pseudomycoides.* Clear zones around the outgrowths of the soil suspensions indicated the presence of at least one strain antagonistic to *B. pseudomycoides* (Fig. 3a). More than 2% of the fast-growing aerobic heterotrophic bacteria suppressed the growth of *B. pseudomycoides* (Fig. 3b). However, the proportion of growth-suppressing bacteria varied widely across the soil samples, with wheatfield soil being the most consistent (Fig. 3b). These findings indicate that soils contain many fast-growing bacteria that are antagonistic to BSL. These results, together with those from native versus sterilized soil microcosms, indicate that the availability of nutrients provided by root exudates benefits other fast-growing bacteria that are antagonistic to BSL, suppressing their population increase.

**Figure 3.**
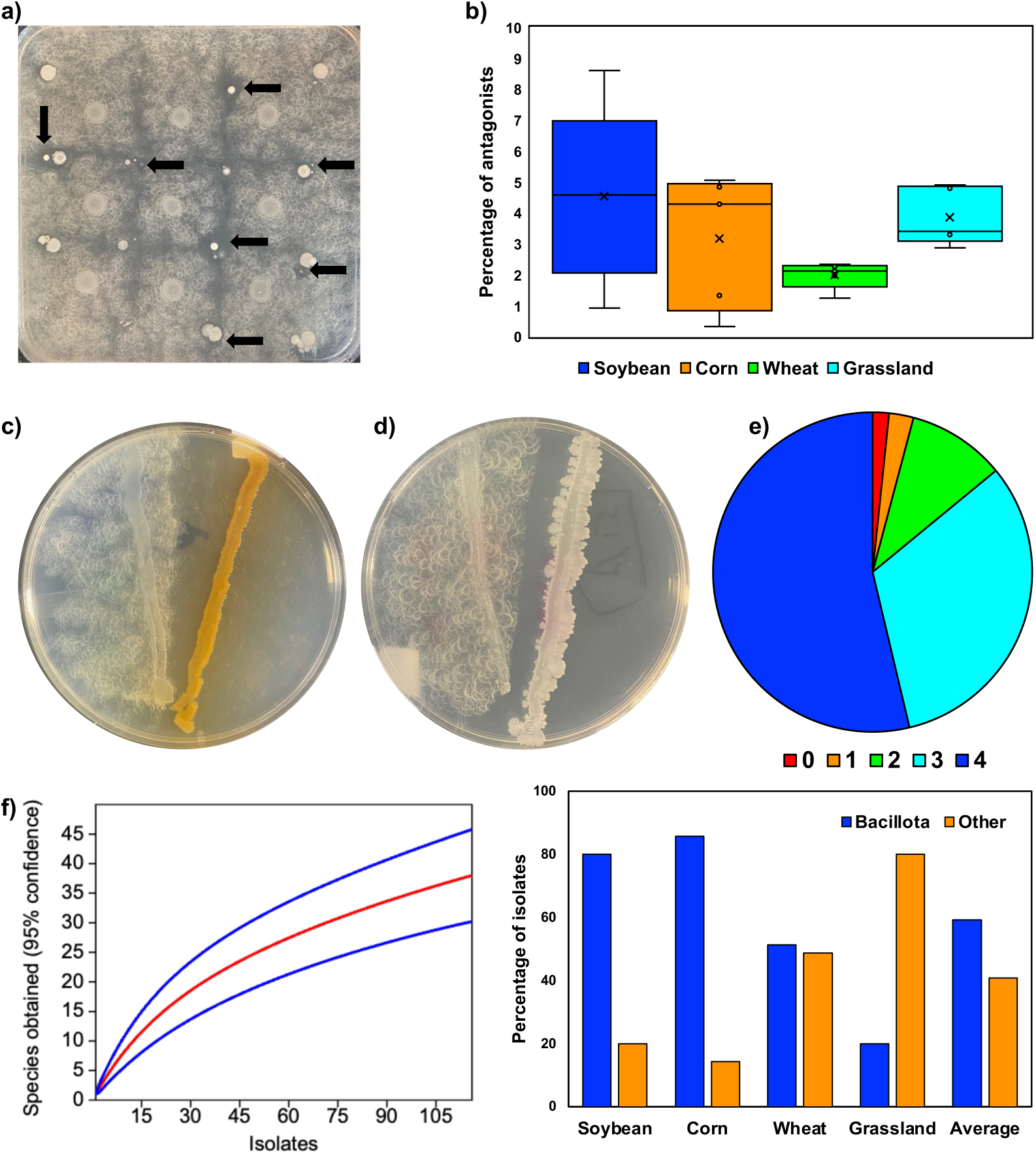
Isolates of rapidly growing aerobic heterotrophs that suppress *Bacillus pseudomycoides.* Serially diluted soil suspensions (10 μL) were spotted between spots of *B. pseudomycoides*, and spots with clear zones indicated the presence of at least one antagonistic strain (a). Percentage of antagonistic bacteria against *B. pseudomycoides* from soil (b). V-streaks of two antagonists plated against *B. pseudomycoides* (c, d). Proportion of isolates that suppressed between 1 and 4 different *Bacillota* (e). Diversity of isolates antagonistic to *B. pseudomycoides,* shown by a rarefaction plot of species richness determined via unconditional variance, with outer lines showing 95% confidence (f), and the proportion of antagonistic *Bacillota* to all other isolates obtained from the four soil types (g).

### Isolation of bacteria antagonistic to BSL

To isolate antagonists of BSL, we overlaid soft agar containing *B. pseudomycoides* over 24 h cultures of soil dilutions on R2A. Zones of inhibition indicated underlying antagonists which were isolated on media that suppress *B. pseudomycoides*. The antagonistic activity of the putative antagonists was confirmed by V-streak (Fig. 3c & d), yielding 124 isolates. To determine whether the isolates also suppressed the growth of other BSL, we performed cross-streak assays against *B. subtilis, B. cereus, P. megaterium* and *P. polymyxa.* The results were striking, as more than half of the isolates exhibited antagonism against all four species (Fig. 3e). Only a small number of antagonists did not demonstrate suppressive activity against any of the BSL. Sequencing of the V1-9 regions of the 16S rRNA genes allowed allocation to the species level in most cases. Most of the 38 species obtained were isolated repeatedly, and rarefaction analysis indicated that this collection is a fair representation of the diversity of fast-growing antagonists of BSL in these soils (Fig. 3f). The percentage of *Pseudomonadota* isolates was 42.7%, and these isolates aligned with *Alfa-*, *Beta-*, and *Gammaproteobacteria* (Fig. S1a). *Actinomycetota* accounted for 2.4% of the population, with one isolate each of *Streptomyces badius, S. camponoticapitis* and *S. lyticus*. The *Bacillota* accounted for 54.8% of the isolates and fell into five genera of the endospore-forming gram-positive rods of the *Bacillaceae* and *Cytobacillaceae* (Fig. S1b). The three monoculture sites contained more BSL than other taxa, whereas antagonists from natural prairie predominantly belonged to *Pseudomonadota* (Fig. 3g). The large number and diversity of BSL isolates found to be antagonistic to *B. pseudomycoides* and to the four other species tested indicated that the majority of BSL antagonists are other BSL, *i.e.*belong to the same physiological group. These results show that soils contain a variety of bacterial taxa that are antagonistic to BSL. These results further indicate that BSL of the *Bacillaceae* and *Cytobacillaceae* play major roles in controlling their collective populations in soil.

### Comparing antagonism among BSL isolates

The large number of *Bacillaceae* and *Cytobacillaceae* bacteria obtained led us to assess interisolate antagonism. Four isolates with the greatest antagonism to *B. pseudomycoides* were selected from each of the seven clades on the BSL phylogenetic tree (Fig. S1b) and cross-streaked. We observed a high number of cases where one isolate suppressed another (Fig. S2, red and blue columns) and a surprising number of cases of mutual growth suppression (Fig. S2, yellow columns). Members of *Brevibacillus*, *Peribacillus* and *B. velezensis* were the most aggressive against other BSL strains, especially against *B. cereus* (Fig. S2). Several pairs of isolates did not inhibit one another (Fig. S2, green columns). The large number of cases of one- and two-way antagonism between pairs of BSL strains indicates that interspecies and strain competitive exclusion are common in this group.

### Genome analysis and prediction of antimicrobial potential

To predict the diversity of antimicrobial compounds produced by BSL isolates antagonistic to other BSL strains, we selected the 10 isolates with the greatest number of antagonistic pairings and the *B. pseudomycoides* tester strain. Genomes annotated via Proka were submitted to antiSMASH to obtain evidence of antimicrobial biosynthesis-related gene clusters (BGCs). We report only clusters that aligned to genes for previously reported antimicrobial compounds, but genomes also contained predicted BGCs with no assigned function (Table S1). While some BGCs aligned well with genes of BSL that produce characterized antimicrobial compounds, others were less related. These findings suggest that the diversity of antimicrobial compounds synthesized by BSL in soil has not been fully characterized. All the isolates contained multiple antimicrobial BGCs (Fig. 4a). *P. megaterium* contained two, whereas *B. velezensis* contained 10 antimicrobial BGCs, while other isolates contained between three and eight. Among them, the eleven genomes encoded 29 different antimicrobial BGCs. The BGCs contained large numbers of genes, with the predicted lengths comprising between 3.4 and 21.1 percent of the genomes (Fig. 4b, Table S1). These findings indicate that antagonism across BSL taxa is mediated by the production of large numbers of different antimicrobial compounds. Species allocation of the 11 sequenced isolates based on whole genome agreed with 16S rRNA gene homolog except for three. Corn66, aligning with *Priestia aryabhattai*, was allocated to the closely related *P. megaterium*, and Grass112 aligning with *Peribacillus simplex* was identified as *P. frigotolerans*. Wheat 6 falling into *B. cereus sensu lato* was identified as *B. tropicus*.

**Figure 4.**
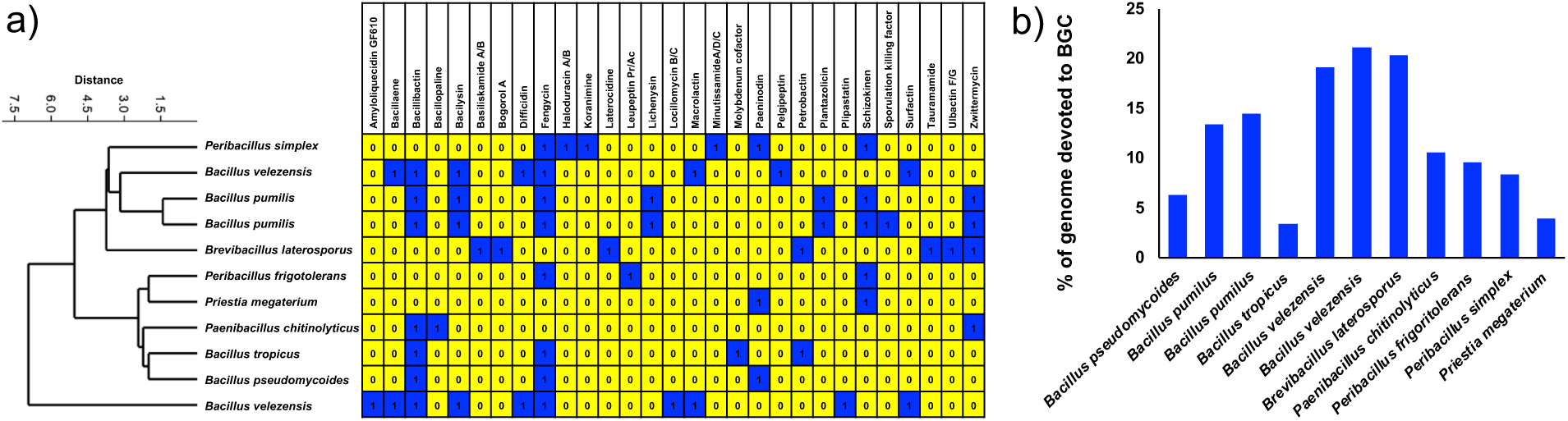
Presence of antimicrobial biosynthesis-related gene clusters (BGC) predicted by antiSmash in the ten sequenced *Caryophanales* genomes and the *B. pseudomycoides* tester strain, with blue indicating presence. The relatedness of BGC profiles was determined using the UPGMA algorithm and Brae Curis similarity index in PAST (a). The percentage of each genome devoted to BGC is shown (b).

### Growth inhibition competition among bacteria

To test whether isolates antagonistic on agar also suppressed the growth of BSL in soil and liquid microcosms, we incubated three gram-negative isolates, one from each of *Alpha-, Beta-,* and *Gammaproteobacteria* (*Bosea robiniae, Pseudomonas chlororaphis*, and *Pseudoduganella guangdongensis*) together with *B. cereus* ATCC 14579, and four gram-positive isolates (*B. cereus*, *B. velezensis, P. megaterium,* and *P. simplex*) separetely. The population density of *B. cereus* alone increased in both the ASM and liquid microcosms, with only some cells forming spores within 48 h (Fig. 5). *B. cereus* did not increase in the presence of the gram-negative consortium in ASM (Fig. 5a & b), indicating growth suppression in soil. In contrast, in the liquid microcosms, there was no support for suppression (Fig. 5c & d). In addition, *B. cereus* sporulated in the presence of the gram-negative consortium in both soil and the liquid (Fig. 5f & h), suggesting that *B. cereus* likely responded to antagonistic pressure by entering sporulation. No population increase was observed in the gram-positive consortia in the soil, indicating mutual suppression among the gram-positive consortium. Sporulation was significantly greater than that in pure cultures in both soil and liquid, indicating the presence of sporulation-inducing pressure among the gram-positive bacteria. The data show that both gram-positive and gram-negative isolates antagonistic on agar were also antagonistic in soil and liquid environments. Collectively, these results support the hypothesis that BSL populations in soil enriched with growth-supporting carbon are suppressed by various other soil bacteria.

**Figure 5.**
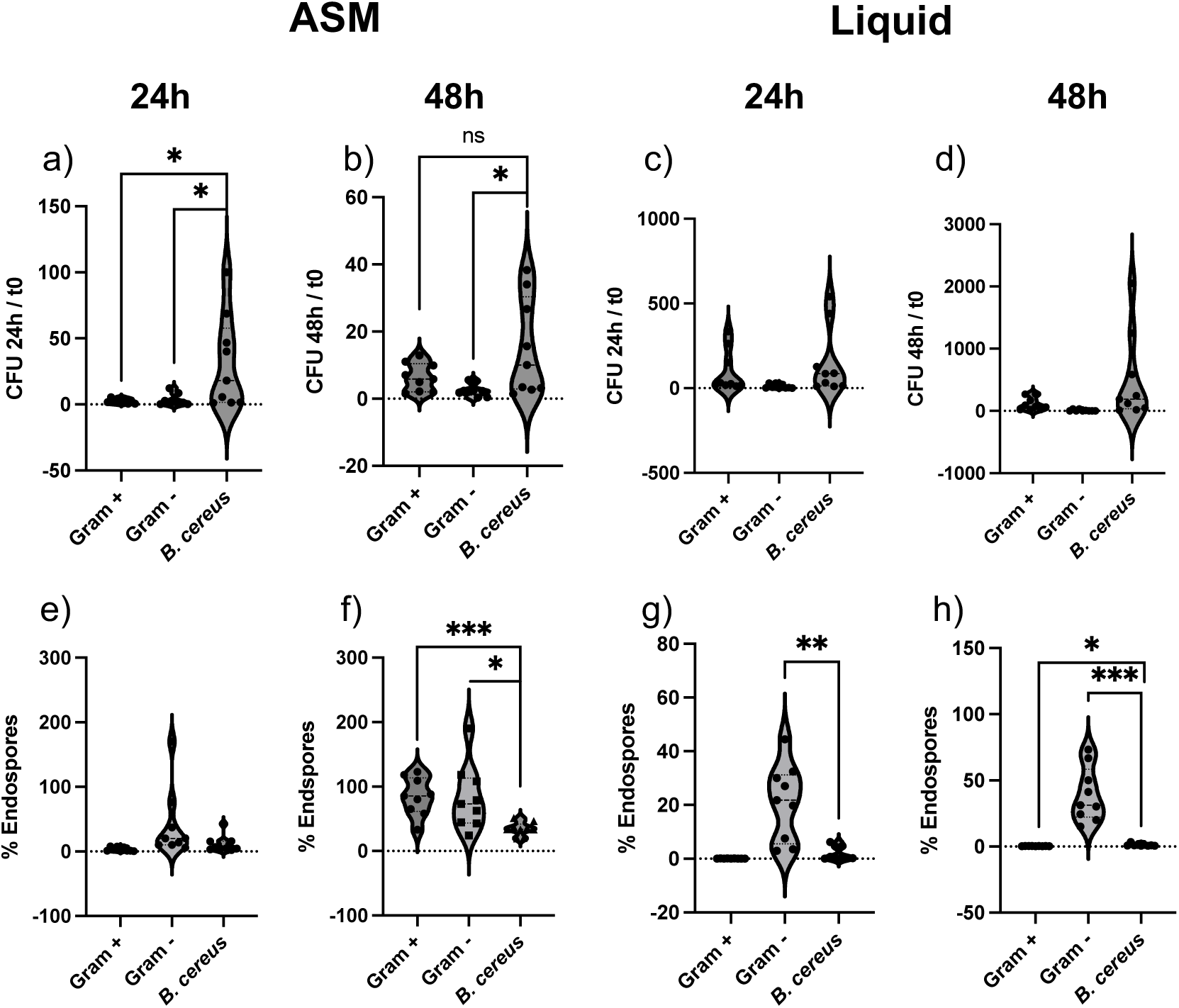
Effect of gram-positive and negative consortia versus *B. cereus* alone after 24 and 48h incubation in either artificial soil microcosms (ASM) or liquid culture supplemented with REC. Violin plots show the ratio of BSL (Gram+ consortium, R2A) and *B. cereus* (Gram-consortium and control, MYP) between t0 and 24 and 48h in ASM (a, b) and liquid (c,d) microcosms. Violin plots show the percentage of endospores of BSL (Gram+ consortium) and *B. cereus* (Gram-consortium and control) between t0 and 24 and 48h in ASM (e, f) and liquid (g,h) microcosms.

Asterisks indicate significant differences between populations according to Welch’s t test (* < 0.05, ** < 0.005, *** < 0.0005). The absence of brackets indicates no significant difference.

## Discussion

We obtained diverse fast-growing bacteria that suppress the growth of *B. pseudomycoides* and other BSL, both on agar and in soil microcosms. These findings support our hypothesis that BSL populations are suppressed by various soil bacteria. The results help explain why these fast-growing copiotrophs, widely viewed as predominant soil bacteria, are less prevalent in the nutrient-rich rhizosphere than in bulk soil. Our culture-independent data show that BSL are less prevalent in the soybean rhizosphere than in bulk soil, and that nutrient amendment does not support an increase in the population of *B. cereus* in the soil environment. These results are supported by previous studies, including meta-analyses of paired clone libraries of bulk soil versus rhizosphere [20], microbiome analysis of the soybean rhizosphere versus bulk soil [24], microbiome analysis of manure-amended versus control soil [25], meta-analyses of multiple paired rhizosphere and bulk soil datasets [22], and our recent study on nutrient-amended wheat field soil [26]. In contrast to population suppression in field soils, *B. cereus* populations increased in nutrient-amended sterilized and synthetic soils, indicating that BSL are hindered in growth by the activities of other soil microbiota. We obtained diverse *Pseudomonadota* and BSL as well as some *Streptomyces* strains that suppressed the population increase in BSL, both on agar and in synthetic soil microcosms. Surprisingly, many of the BSL isolates inhibited each other, suggesting complex competitive relationships within the BSL group. The results presented here support our hypothesis that BSL are suppressed by other antagonistic bacteria in soil.

### Restraints on the population increase of BSL

The population dynamics of BSL are unique among bacteria because a decrease in vegetative cell numbers could be due to either death or sporulation. Likewise, an increase in the vegetative count could be due either to cell division (growth) or to germination of endospores. Our results indicate that the population restraint of BSL in response to other microbiota is due to the suppression of cell division and of germination, and to sporulation. Our previous study also showed similar results: *B. cereus sensu lato* did not increase with the availability of nutrients [26]. The inhibition of colony expansion on agar inhibits population growth but cannot distinguish between growth suppression and sporulation. The growth of BSL in the soil microcosms was suppressed in the presence of other BSL strains, including *B. cereus* and *B. velezensis* (Fig. 5a, e). *B. velezensis* produces different antimicrobial agents reported to suppress *B. cereus* [65]. *B. cereus* was inhibited by three other *Bacillus* species in the rhizosphere of Bambara groundnut [66]. The growth of the BSL consortium was not fully suppressed by other BSL strains in the liquid microcosms, and the degree of sporulation was very low. This may be because the mechanism of antagonism requires closer proximity to suppress BSL growth. *B. cereus* was also suppressed by the gram-negative consortium in both ASM and liquid media (Fig. 5 a–d). The consortium included a *Burkholderia* isolate, and *Burkholderia thailandensis* reportedly produces various antibiotics, including bactobolin variant A (thailandene A), which has shown cryptic antibacterial activity to inhibit *B. subtilis* [67]. Interestingly, a large proportion of the cells sporulated in the presence of gram-negative-negative bacteria, as indicated by the high percentage of spores that formed (Fig. 5 e–h).

Within 48 h of exposure to the gram-positive and gram-negative consortia in the soil, the BSL were largely in spore form, whereas the *B. cereus* single culture contained mostly vegetative cells paired with a population increase (Fig. 5f, b). *B. cereus* is known to sporulate profusely in the absence of external pressure (Fig. 5) [68], but the proportion of sporulated cells was far greater when it was exposed to other species (Fig. 5f). Importantly, the introduction of *B. cereus* into sterile soybean soil led to ∼100% sporulation, whereas in ASM, only ∼1% of the cells formed spores (Fig. 2f, g). As ASM never contained any bacteria, sporulation in sterilized soybean soil was possibly because of heat-stable products of the resident microbiota. Several gram-negative bacteria have been reported to induce sporulation in *B. subtilis* by activating the histidine kinases KinA and KinB, which are key regulators of sporulation in *B. subtilis*. These inducers include *Escherichia coli* [69], *Salmonella typhimurium* [70] and *P. chlororaphis* [71]. Overall, the gram-negative antagonists appeared to suppress BSL population growth in large part by inducing sporulation. While halting population growth, the resulting endospores can support future population expansion through germination.

Our data revealed that the soybean and wheat soils contained BSL endospores that did not germinate, as the spore numbers remained unchanged with increasing nutrient availability (Fig. 2b, c). While some vegetative cells were present in corn and natural grasslands, the spore number also did not decrease (Fig. 2d, e). These spores were enumerated as colonies formed on MYP after heat treatment, showing that they were still viable and able to germinate once placed in an environment free from other soil microbiota. Thus, BSL spore germination likely was suppressed in native soils amended with REC. The vegetative outgrowth in sterile soybean soil and ASM following the addition of REC showed that some of the endospores germinated (Fig. 2 f, g). Far less is known about cues for germination than spore formation, but *B. subtilis* and *B. cereus* germination receptors are responsive to nutrients such as L-alanine and L-glutamine [72, 73]. While the activation of germination receptors initiates the process of germination, several compounds, such as nisin (*Lactococcus lactis*) and thusin (*B. thuringiensis*), inhibit the outgrowth of spores primed for germination [74, 75]. Germinated endospores inhibited in outgrowth are no longer viable, as they have lost the resilience of endospores and are unable to grow into full cells. The high numbers of endospores detected from several microcosms by culturing were, therefore, more likely subject to repression of germination because the REC included alanine and glycine, both of which are known to activate germination. The nature and mechanisms underlying repressors of endospore germination in soils are only cursorily known [76] and require urgent attention.

### Bacterial groups contributing to the population restraint of BSL

The observed population restraint of BSL was caused by various bacterial groups in *Pseudomonadota*, some *Actinomycetota*, and, surprisingly, many BSL. This aligns well with a broader understanding of the rhizosphere, where nutrient provision supports the growth of diverse copiotrophs, some of which are involved in chemical warfare. *Pseudomonadota* includes many bacteria known for producing antibacterial compounds [77, 78]. For example, *Pseudomonas* are known for producing antimicrobials such as phenazines, siderophores, and lipopeptides that target both fungal and bacterial rivals [79, 80]. *Pseudomonas* are known to specifically suppress various *Bacillus* [38, 81]. *Streptomyces* are well-known producers of antibiotics such as streptomycin [34]. In addition to diverse antibiotics, some *Streptomyces* produce siderophores that sensitize *Bacillus* to phage infection by inhibiting the activation of Spo0A, a key regulator of sporulation [82]. The vegetative cells may be vulnerable to attack by other antimicrobial agents.

The most striking finding from our study was the broad antagonism among members of BSL. Many of these were isolated from the same ecological niches. Many BSL species are able to produce antimicrobial compounds such as lipopeptides that suppress other microbiota, such as plant pathogens. Under biculture conditions, these copiotrophic BSL tend to outcompete other microbes in the rhizosphere well, supporting their status as biological control agents [83, 84]. There are multiple recent reports of BSL isolates that suppress the growth of select others. *B. velezensis* secretes LXG toxins via a type VII secretion system to suppress *B. proteolyticus* and other strains of *B. velezensis* in biofilms [85]. A strain of *B. velezensis* was found to suppress *B. subtilis* in the mouse gut [86]. *B. amyloliquefaciens*, *B. thuringiensis*, and an unidentified *Bacillus* obtained from the Bambara groundnut rhizosphere produced butanol-extracted volatile organic compounds that suppress the growth of *B. cereus* [66]. Under nutrient limitation, *B. subtilis* produces cannibalistic toxins that selectively kill neighboring nontoxin-producing isogenic siblings, enabling the surviving cells to recycle nutrients and delay sporulation [87]. Our results indicate that BSL antimicrobials also act against many other BSL strains, directing defense mechanisms in closely related taxa. The onset of sporulation is inhibited by some microbial products that inhibit the phosphorylation of Spo0A. *B. velezensis* produces the quorum-sensing molecule autoinducer-2, which inhibits the activation of its own Spo0A [88]. The cells remaining in vegetative form are vulnerable to attack by other antimicrobial agents. Conversely, our data indicate that BSL are competitive even among closely related taxa because of the production of diverse antimicrobial compounds or inducers of sporulation. In the nutrient-rich and highly competitive rhizosphere, BSL species appear to have evolved a strategy of aggression over their coexistence such that almost every strain of BSL is a rival. This is different from *Streptomyces,* where there is little to no competition among close relatives [89].

### Diversity of biosynthetic gene clusters

BGCs constituted between 3.4 and 21.1% of the BSL genomes reported here, and each encoded between 2 and 10 BGCs, with 29 different BGCs overall. This is consistent with findings from large-scale genome mining studies, where BSL typically carry BGCs, including those for nonribosomal peptide synthetases (NRPs), polyketide synthases (PKSs), and ribosomally synthesized peptides (RiPPs) [90]. A recent study of ten sequenced *Bacillus* strains reported 120 BGCs belonging to the NRPSs surfactin (8 BGCs), fengycin (8 BGCs), bacillibactin (10 BGCs), PKS macrolactin (2 BGCs), difficidin (2 BGCs), hybrid NRPS/PKS bacillaene (8 BGCs), bacteriocin subtilin (2 BGCs), and subtilosin A (6 BGCs) [91]. *B. pumilus* SF-4 produces 12 BGCs [92]. This indicates a major genomic investment in encoding antimicrobial compounds and a common strategy for competition. The encoding and synthesis of these compounds may be for defensive or offensive reasons, but both come at a substantial metabolic cost to each cell, restricting its fitness. Closely related BSL strains tended to carry overlapping but distinct sets of BGCs (Fig. 4), supporting the antagonism observed among members of the same species (Fig. S2). Importantly, this approach appears to be common across BSL, so despite its highly effective copiotrophic traits, the group appears to limit itself in soil and the rhizosphere. While the rapid growth rate and metabolic traits of BSL make it an invaluable workhorse in biotechnology [93] and in the biocontrol and growth promotion of crop plants [94], our data indicate that it could be vulnerable to the surrounding microbiota in soils.

### Ecophysiology of BSL in soil revisited

In 1916, Joel Conn asked, “Are spore-forming bacteria of any significance in soil under normal conditions?” [5], concluding that most *Bacillus* in soil occur as endospores and do not contribute to activity in the soil. The proportion of BSL in soil that occur as spores versus vegetative cells has not been well established. However, our data show that BSL are either induced to sporulate or inhibited from germination by diverse soil bacteria, including *Pseudomonadota* and BSL. This limits their activity in soil to niches where they are protected from other copiotrophs or where their arsenal of biosynthetic genes encodes compounds that suppress other bacteria. Whether BSL species grow, persist in the stationary phase or persist as endospores in soil and in the rhizosphere is a critical question for microbial ecology.

## Data availability

16S rRNA data available here: https://www.ncbi.nlm.nih.gov/nuccore/?term=PV889183:PV889306[accn]

Whole-genome data available here: Awaiting release by NCBI

## Acknowledgments

Muhammad Yasir Afzal was supported by the Higher Education Commission of Pakistan (HEC-Pakistan) for PhD scholarship programs under the “US-Pakistan Knowledge Corridor”. We thank Prof Fanus Venter for helpful discussions and for proposing the term *Bacillus sensu lato*. This material is based upon work conducted via the South Dakota State University Genomics Sequencing Core Facility (RRID:SCR_023959), which is supported in part by the South Dakota Agricultural Experiment Station, and by the National Institute of General Medical Sciences of the National Institutes of Health under Award Number P20GM135008. The content is solely the responsibility of the authors and does not necessarily represent the official views of the National Institutes of Health.

## Funding

This research was funded by the South Dakota Agricultural Experiment Station.

## Author contributions

MYA, and VB designed the study with inputs from JK. VB planned the experiments and supervised the project. MYA and JK performed experiments. MYA, JK and VB analyzed the data. MYA prepared the original draft and MYA, JK and VB revised the manuscript.

## Notes

### Competing Interest Statement

The authors have declared no competing interest.

