## Supplementary material for "Fast-growing *Bacillus sensu lato* rhizosphere populations are constrained by antagonistic Pseudomonadota, Actinomycetota *and other* Bacillus sensu lato": Fig. S1

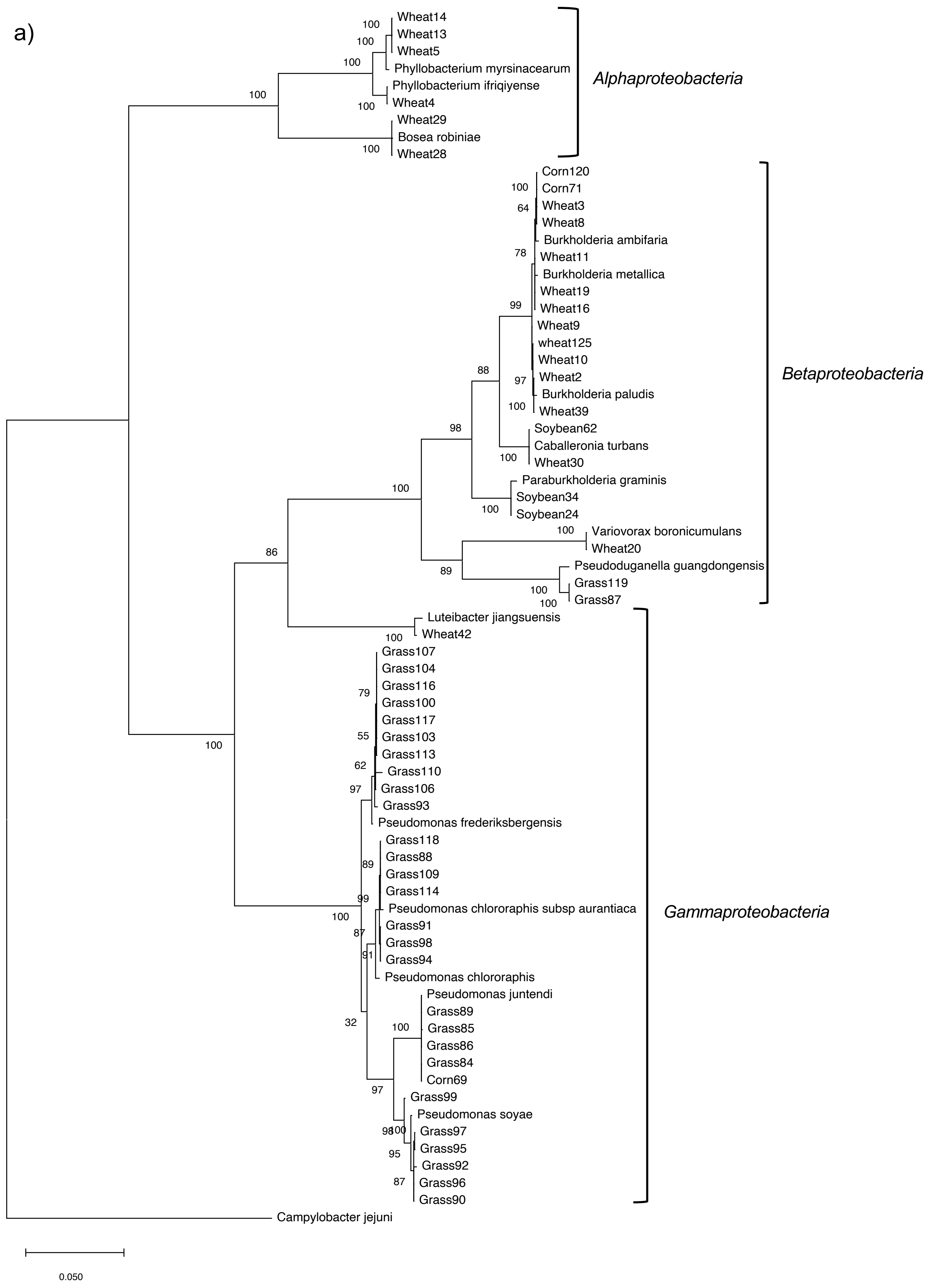


Figure S1. Diversity of antagonists of *B. pseudomycoides* obtained. The diversity of *Pseudomonadota* (a) and *Bacillota* (b) isolates obtained is shown by maximum likelihood of the V1-9 region of the 16SrRNA genes with 100 bootstraps.

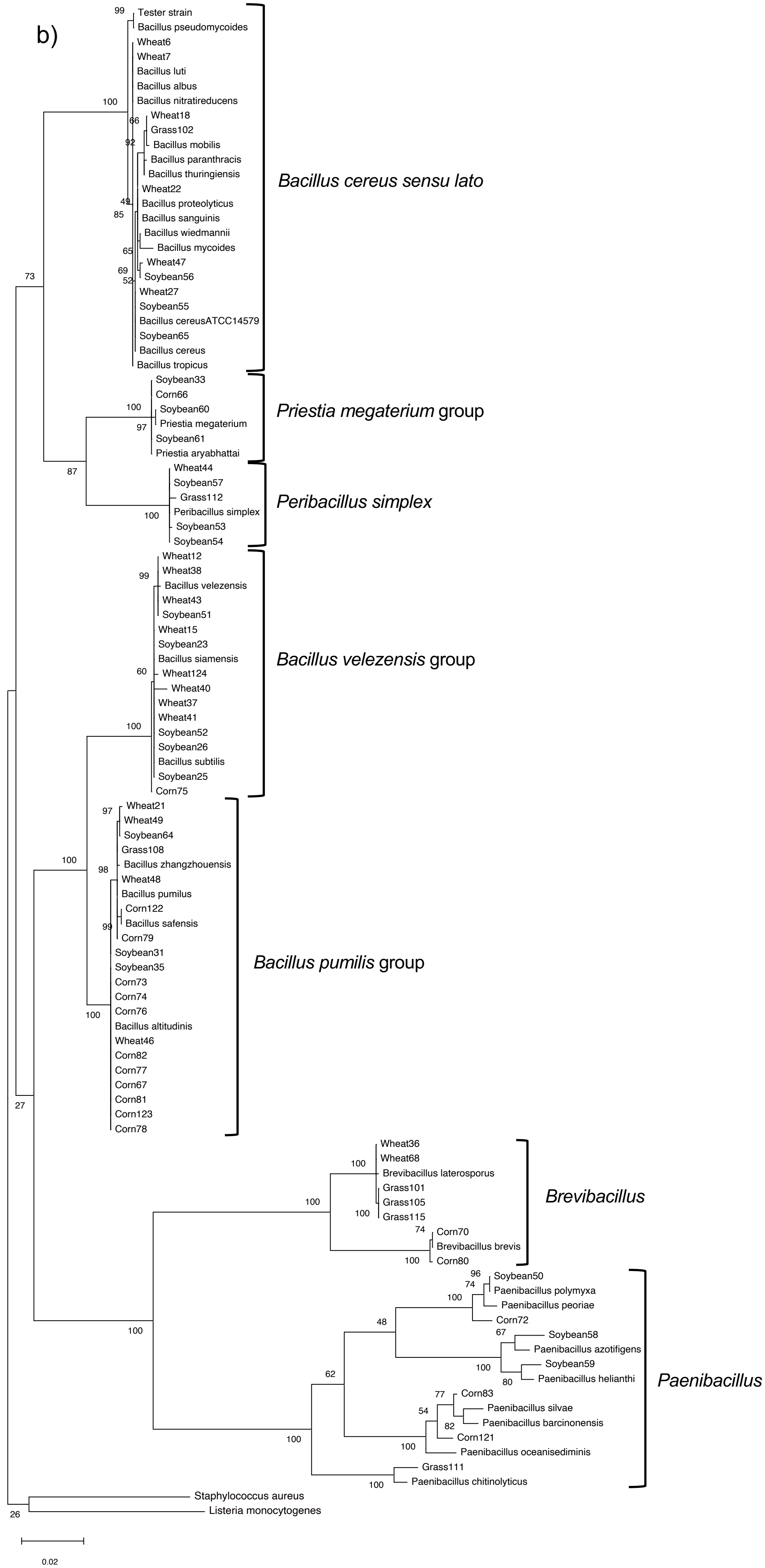
b)
Tester strain
Bacillus pseudomycoides
Wheat6
Wheat7
Bacillus luti
Bacillus albus
Bacillus nitratireducens
Wheat18
Grass102
Bacillus mobilis
Bacillus paranthracis
Bacillus thuringiensis
Wheat22
Bacillus proteolyticus
Bacillus sanguinis
Bacillus wiedmannii
Bacillus mycoides
Wheat47
Soybean56
Wheat27
Soybean55
Bacillus cereusATCC14579
Soybean65
Bacillus cereus
Bacillus tropicus
*Bacillus cereus sensu lato*
Soybean33
Corn66
Soybean60
Priestia megaterium
Soybean61
Priestia aryabhattai
*Priestia megaterium* group
Wheat44
Soybean57
Grass112
Peribacillus simplex
Soybean53
Soybean54
*Peribacillus simplex*
Wheat12
Wheat38
Bacillus velezensis
Wheat43
Soybean51
Wheat15
Soybean23
Bacillus siamensis
Wheat124
Wheat40
Wheat37
Wheat41
Soybean52
Soybean26
Bacillus subtilis
Soybean25
Corn75
*Bacillus velezensis* group
Wheat21
Wheat49
Soybean64
Grass108
Bacillus zhangzhouensis
Wheat48
Bacillus pumilus
Corn122
Bacillus safensis
Corn79
Soybean31
Soybean35
Corn73
Corn74
Corn76
Bacillus altitudinis
Wheat46
Corn82
Corn77
Corn67
Corn81
Corn123
Corn78
*Bacillus pumilis* group
Wheat36
Wheat68
Brevibacillus laterosporus
Grass101
Grass105
Grass115
Corn70
Brevibacillus brevis
Corn80
*Brevibacillus*
Soybean50
Paenibacillus polymyxa
Paenibacillus peoriae
Corn72
Soybean58
Paenibacillus azotifigens
Soybean59
Paenibacillus helianthi
Corn83
Paenibacillus silvae
Paenibacillus barcinonensis
Corn121
Paenibacillus oceanisediminis
Grass111
Paenibacillus chitinolyticus
*Paenibacillus*
Staphylococcus aureus
Listeria monocytogenes
99
100
66
92
49
85
65
69
52
73
100
97
87
100
99
60
100
100
97
98
99
100
27
100
100
100
74
100
96
74
100
48
67
100
80
62
77
54
82
100
100
100
100
26
0.02
