## Supplementary material for "Fast-growing *Bacillus sensu lato* rhizosphere populations are constrained by antagonistic Pseudomonadota, Actinomycetota *and other* Bacillus sensu lato": Fig. S2

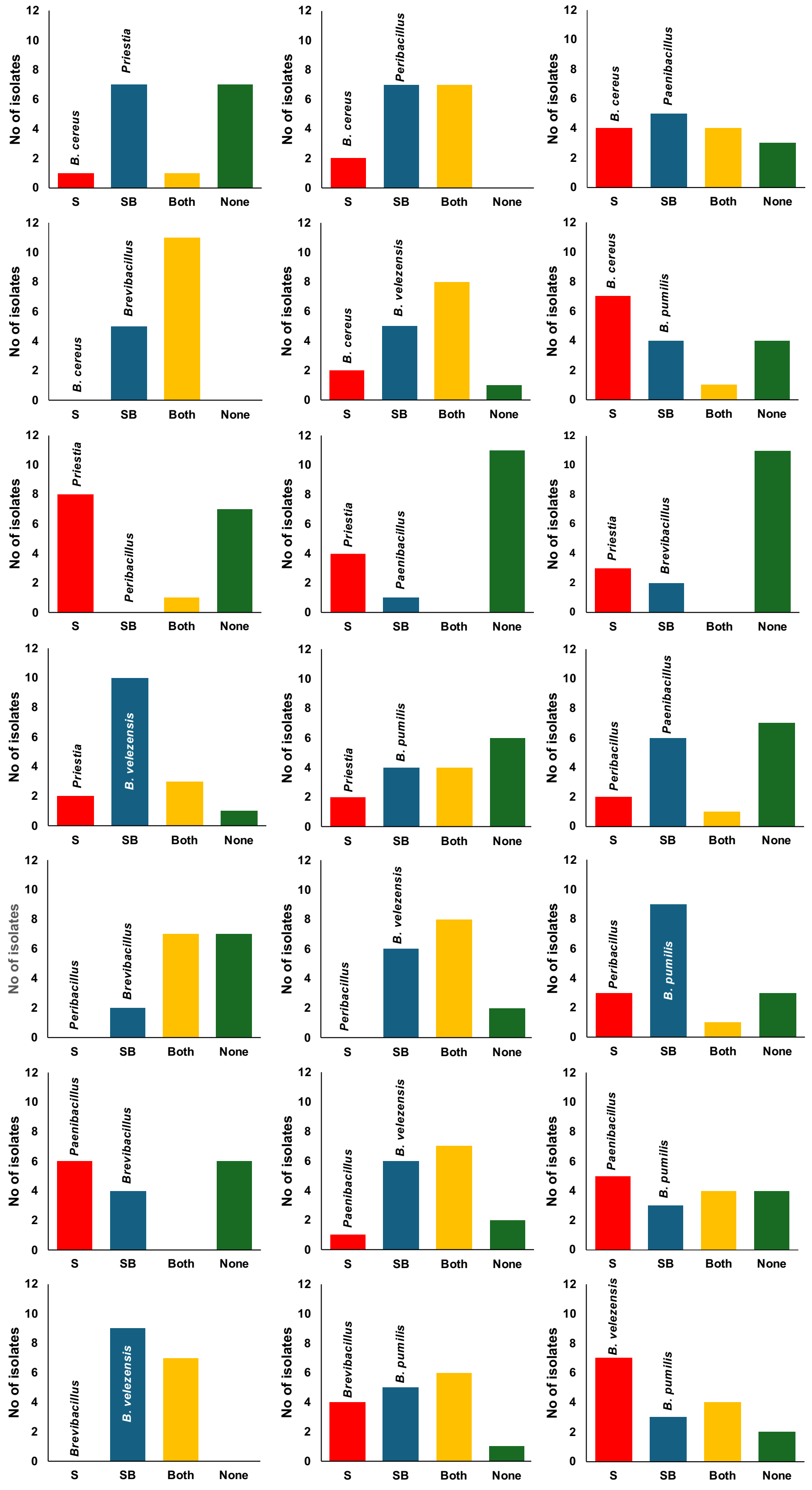

Figure S2. Antagonism among representatives of the *Bacillus* antagonistic isolates. Four isolates from each of the seven clades were cross-streaked to determine suppression of other taxa (S, red), suppression by other taxa (SB, blue), two-way suppression (Both, yellow) and absence of suppression (None, green).
