## Supplementary material for "Fast-growing *Bacillus sensu lato* rhizosphere populations are constrained by antagonistic Pseudomonadota, Actinomycetota *and other* Bacillus sensu lato": Table S1

Table S.1 BGC's in the 11 genomes of BSL as identified using antiSMASH.

| Isolate ID and Name | Region | Type | Genome (BPs) From | To | Genome investment (BPs) | Most similar known cluster | Similarity | Cluster similirty with bacterial species | %age Cluster similirty with bacterial species |
|---|---|---|---|---|---|---|---|---|---|
| | 1 | lassopeptide | 317,687 | 341,604 | 23917 | Paeninodin | | | 100% of genes show similarity |
| **TBC_tester_strain_wheat1_sample1** | 2 | terpene like precursor | 1,303,373 | 1,324,266 | 20893 | | | | |
| ***Bacillus pseudomycoides*** | 3 | terpene | 1946115 | 1967974 | 21859 | | | | |
| | 4 | RIPP-like | 2664962 | 2676836 | 11874 | | | | |
| **Genome size: 5121675 BPs** | 5 | Betalactone | 2708520 | 2733757 | 25237 | Fengycin | | | 40% of genes show similarity |
| | 6 | NRP metallophore | 2807304 | 2914351 | 107047 | bacillibactin | | | 85% of gennes show similarity |
| | 7 | NRPS | 3557804 | 3620984 | 63180 | | | | |
| | 8 | ranthipeptide | 3898683 | 3935675 | 36992 | | | | |
| | 9 | RIPP-like | 4190509 | 4201747 | 11238 | | | | |
| **Total genome investment for toxin production** | | | | | 322237 | | | | |
| | 1 | Azole-containing-RIPP | 909066 | 932572 | 23506 | | | | |
| **Wheat 6** | 2 | Ni-siderophore | 1559477 | 1591183 | 31706 | Petrobactin | 100% | Bacillus cereus strain AR156 | 100% of genes show similarity |
| ***Bacillus tropicus*** | 3 | NRP-metallophore, NRPS | 1873452 | 1925199 | 51747 | Bacillibactin | 85% | Bacillus tropicus strain AOA-CPS1 | 100% of genes show similarity |
| | 4 | Betalactone | 2086228 | 2111466 | 25238 | Fengycin | 40% | Bacillus cereus strain MOD1 Bc216 Bc216 | 100% of genes show similarity |
| **Genome size: 5145247 BPs** | 5 | RIPP-like | 2158328 | 2168627 | 10299 | | | | |
| | 6 | RIPP-like | 2214846 | 2225112 | 10266 | | | | |
| | 7 | Terpene | 3015617 | 3037470 | 21853 | Molybdenum cofactor | 17% | Bacillus cereus strain AR156 | 100% of genes show similarity |
| | 8 | Terpene | 3631840 | 3652730 | 20890 | | | | |
| **Total genome investment for toxin production** | | | | | 174615 | | | | |
| | 1 | NRPS | 1 | 29545 | 29544 | Surfactin | 52% | Bacillus velezensis YAU B9601-Y2 | 46% of genes show similarity |
| **Soybean 23** | 2 | Other | 636123 | 677541 | 41418 | Bacilysin | 100% | Bacillus velezensis strain BS-37 chromosome | 100% of genes show similarity |
| ***Bacillus velezensis*** | 3 | NRPS | 878359 | 946145 | 67786 | Pelgipeptin | 25% | Bacillus amyloliquefaciens strain JP3042 | 100% of genes show similarity |
| | 4 | RIPP like, NRP mettalophore, NRPS | 1293341 | 1344185 | 50844 | Bacillibactin | 100% | Bacillus velezensis strain BIM B-439D | 97% of genes show similarity |
| **Genome size: 4029617 BPs** | 5 | Terpene | 1961067 | 1981957 | 20890 | | | | |
| | 6 | transAT-PKS | 2021635 | 2127804 | 106169 | Difficidin | 100% | Bacillus velezensis strain LBUM279 | 95% of genes show similarity |
| | 7 | T3PKS | 2243367 | 2284467 | 41100 | | | | |
| | 8 | Terpene | 2334844 | 2356766 | 21922 | | | | |
| | 9 | NRPS Betalctone NRPS like | 2380542 | 2517630 | 137088 | Fengycin | 100% | Bacillus velezensis strain BIOMA BV10 | 96% of genes show similarity |
| | 10 | transAT-PKS NRPS like NRPS T3PKS | 2579019 | 2689120 | 110101 | Bacillaene | 100% | Bacillus velezensis strain BS-37 | 100% of genes show similarity |
| | 11 | transAT-PKS | 2911267 | 2999481 | 88214 | Macrolactin | 100% | Bacillus velezensis strain S4 | 95% of genes show similarity |
| | 12 | Terpene | 3301371 | 3322111 | 20740 | | | | |
| | 13 | NRPS | 3993772 | 4029618 | 35846 | Surfactin | 47% | Bacillus velezensis strain LBUM279 | 57% of genes show similarity |
| **Total genome investment for toxin production** | | | | | 771662 | | | | |
| | 1 | T1PKS NRPS | 211286 | 290439 | 79153 | Zwittermycin | 18% | Bacillus pumilus strain NRRL BD-222 | 95% of genes show similarity |
| **Wheat 49** | 2 | NRPS | 509989 | 593357 | 83368 | Lichenysin | 85% | Bacillus pumilus strain NRRL B-59349 | 100% of genes show similarity |
| **Bacillus pumilus** | 3 | NRP mettalophoe NRPS | 959989 | 1010985 | 50996 | Bacillibactin Bacillibactin E, Bacillibactin F | 80% | Bacillus pumilus strain BIM B-171 | 100% of genes show similarity |
| | 4 | RIPP like | 1063157 | 1073501 | 10344 | | | | |
| **Genome size: 3838066 BPs** | 5 | Other | 1240693 | 1282114 | 41421 | Bacilysin | 85% | Bacillus pumilus strain NRRL NRS-0334 | 97% of genes show similarity |
| | 6 | Terpene | 1827153 | 1848076 | 20923 | | | | |
| | 7 | Betalactone | 2176150 | 2208454 | 32304 | | | | |
| | 8 | T3PKS | 2703173 | 2744273 | 41100 | | | | |
| | 9 | Terpene | 2782368 | 2804242 | 21874 | | | | |
| | 10 | Betalactone | 2867254 | 2895670 | 28416 | Fengycin | 53% | Bacillus pumilus strain Ha06YP001 | 100% of genes show similarity |
| | 11 | Terpene | 2782368 | 2804242 | 21874 | | | | |
| | 12 | LAP RRE containing | 3096436 | 3119596 | 23160 | Plantazolicin | 91% | Bacillus safensis strain BRM1 | 95% of genes show similarity |
| | 13 | NI siderophore / terpene | 3626323 | 3663961 | 37638 | Schizokinen | 60% | Bacillus safensis strain NRRL B-03275 | 100% of genes show similarity |
| | 14 | RRE containing | 3813236 | 3834141 | 20905 | | | | |
| **Total genome investment for toxin production** | | | | | 513476 | | | | |
| | 1 | NI siderophore | 429617 | 463130 | 33513 | Schizokinen | 75% | Peribacillus simplex strain SH-B26 | 92% of genes show similarity |
| **Soybean 53** | 2 | Cyclic lactone autoinducer RIPP like lanthipeptide clss ii | 701116 | 736746 | 35630 | HaloduracinB / haloduricina | 40% | Ureibacillus thermosphaericus strain A1 | 23% of genes show similarity |
| ***Peribacillus simplex*** | 3 | Terpene | 800471 | 822432 | 21961 | CitillinA / Citillin B | 7% | Peribacillus simplex strain SH-B26 | 100% of genes show similarity |
| | 4 | T1PKS | 1282374 | 1324914 | 42540 | MinutissamideA / D/ C | 23% | Peribacillus frigoritolerans strain KF19 | 33% of genes show similarity |
| **Genome size: 5393512 BPs** | 5 | Azole-containing-RIPP | 1403356 | 1426894 | 23538 | | | | |
| | 6 | Lassopeptide | 2100882 | 2124851 | 23969 | Paeninodin | 80% | Peribacillus frigoritolerans strain NS1 | 80% of genes show similarity |
| | 7 | Terpene | 2289901 | 2310794 | 20893 | | | | |
| | 8 | Terpene | 2333740 | 2354558 | 20818 | | | | |
| | 9 | T3PKS | 2492581 | 2533669 | 41088 | | | | |
| | 10 | Cyclic lactone autoindicer NRPS | 3304536 | 3352018 | 47482 | | | | |
| | 11 | Betalactone NRPS | 3359267 | 3438236 | 78969 | Fengycin | 46% | Peribacillus simplex strain SH-B26 | 86% of genes show similarity |
| | 12 | NRPS | 5000267 | 5060574 | 60307 | Koranimine | 87% | Peribacillus simplex strain SH-B26 | 85% of genes show similarity |
| **Total genome investment for toxin production** | | | | | 450708 | | | | |
| | 2 | Terpene | 329801 | 351669 | 21868 | | | | |
| **Corn 66** | 3 | Terpene | 1902149 | 1922967 | 20818 | Surfactin | 13% | Priestia megaterium strain AFS004604 | 100% of genes show similarity |
| ***Priestia megaterium*** | 4 | NI-siderophore | 2073478 | 2108053 | 34575 | Schizokinen | 62% | Priestia megaterium strain SGAir0080 | 100% of genes show similarity |
| | | Terpene | 2478633 | 2499529 | 20896 | | | | |
| **Genome size: 5096019 BPs** | 5 | Terpene | 3909736 | 3930584 | 20848 | Carotenoid | 50% | Priestia megaterium strain AFS004604 | 100% of genes show similarity |
| | 6 | Phosphonate | 4054437 | 4071859 | 17422 | | | | |
| | 7 | Lasspeptide | 4374752 | 4398660 | 23908 | Paeninodin | 100% | Priestia megaterium strain F16 | 76% of genes show similarity |
| | 8 | T3PKS | 4713043 | 4754128 | 41085 | | | | |
| **Total genome investment for toxin production** | | | | | 201420 | | | | |

### Table S.1 BGC’s in the 11 genomes of BSL as identified using antiSMASH.

| | | | | | | | | | |
|---|---|---|---|---|---|---|---|---|---|
| | 1.1 | NRPS | 1 | 50264 | 50263 | Plipastatin | 53% | Bacillus velezensis strain MBNC DBT-NECAB | 30% of genes show similarity |
| **Corn 75** | 1.2 | Terpene | 75541 | 97424 | 21883 | | | | |
| **Bacillus velezensis siamensis group** | 1.3 | T3PKS | 164207 | 205307 | 41100 | | | | |
| | 1.4 | transAT-PKS | 320174 | 426342 | 106168 | Difficidin | 100% | Bacillus velezensis strain FJAT-46737 | 96% of genes show similarity |
| **Genome size: 4019 Kbs** | 1.5 | NRP mettalophore NRPS Ripp lik | 1116894 | 1168360 | 51466 | Bacillibactin | 100% | Bacillus velezensis strain LBUM279 | 97% of genes show similarity |
| | 1.6 | Other | 1694973 | 1736391 | 41418 | Bacilysin | 100% | Bacillus velezensis strain SW5 | 100% of genes show similarity |
| | 1.7 | NRPS transAT-PKS | 2230788 | 2308523 | 77735 | Locillomycin B/C | 28% | Bacillus velezensis strain LBUM279 | 100% of genes show similarity |
| | 1.8 | NRPS transAT-PKS | 2387614 | 2453018 | 65404 | Surfactin | 91% | Bacillus velezensis strain LBUM279 | 97% of genes show similarity |
| | 1.9 | Cyclic lactone autoinducer Lnthi | 2663031 | 2692446 | 29415 | Amyloliquecidin GF610 | 93% | Bacillus velezensis strain LBUM279 | 100% of genes show similarity |
| | 1.10 | PKS-like | 3015977 | 3057221 | 41244 | Butirosin A/B | 7% | Bacillus velezensis YAU B9601-Y2 | 100% of genes show similarity |
| | 1.11 | Terpene | 3140009 | 3160749 | 20740 | | | | |
| | 1.12 | transAT-PKS | 3538249 | 3626333 | 88084 | Macrolactin H | 100% | Bacillus velezensis strain SW5 | 91% of genes show similarity |
| | 1.13 | transAT-PKS T3PKS NRPS | 3850313 | 3960350 | 110037 | Bacillaene | 100% | Bacillus velezensis strain 9D-6 | 96% of genes show similarity |
| | 1.14 | NRPS transAT-PKS betalactone | 4021287 | 4108817 | 87530 | Fengycin | 73% | Bacillus velezensis strain VTX9 | 73% of genes show similarity |
| **Total genome investment for toxin production** | | | | | 832487 | | | | |
| | 1.1 | NRPS T1PKS | 68098 | 149328 | 81230 | Zwittermycin | 18% | Bacillus pumilus strain Ha06YP001 | 100% of genes show similarity |
| **Corn 79** | 1.2 | RRE containing | 324999 | 345904 | 20905 | | | | |
| **Bacillus pumilus** | 1.3 | Terpene NI-siderophore | 487815 | 525471 | 37656 | Schizokinen | 60% | Bacillus safensis strain 1370ba1 | 100% of genes show similarity |
| | 1.4 | RRE-containing LAP | 1014407 | 1037570 | 23163 | Plantazolicin | 91% | Bacillus safensis strain BRM1 | 95% of genes show similarity |
| **Genome size: 3644 Kbs** | 1.5 | Betalactone | 1217426 | 1245844 | 28418 | Fengycin | 53% | Bacillus pumilus strain Ha06YP001 | 100% of genes show similarity |
| | 1.6 | Terpene | 1313843 | 1335717 | 21874 | | | | |
| | 1.7 | T3PKS | 1374029 | 1415129 | 41100 | | | | |
| | 1.8 | Betalactone | 1910077 | 1942384 | 32307 | | | | |
| | 1.9 | Other | 2786270 | 2827691 | 41421 | Bacilysin | 85% | Bacillus pumilus strain NRRL NRS-0334 | 93% of genes show similarity |
| | 1.10 | NBRP=metallophore NRPS | 3057909 | 3109635 | 51726 | Bacillibacton E/F | 80% | Bacillus pumilus strain NRRL B-59349 | 100% of genes show similarity |
| | 1.11 | Sactipeptide ranthipeptide RIPP | 3311861 | 3344679 | 32818 | Sporulation killing factor | 85% | Bacillus pumilus strain UAMX | 100% of genes show similarity |
| | 1.12 | NRPS | 3481537 | 3565235 | 83698 | Lichenysin | 85% | Bacillus pumilus strain BIM B-171 c | 100% of genes show similarity |
| **Total genome investment for toxin production** | | | | | 496316 | | | | |
| | 1 | RRE containing | 236452 | 257612 | 21160 | Pf-5 pyoverdine | 1% | Brevibacillus laterosporus DSM 25 | 100% of genes show similarity |
| **Grass 105** | 2 | Cyclic lactone autoinducer | 623594 | 644109 | 20515 | | | | |
| **Brevibacillus laterosporus** | 3 | NI-siderophore | 766540 | 798246 | 31706 | Petrobactin | 100% | Brevibacillus laterosporus DSM 25 | 100% of genes show similarity |
| | 4 | NRPS lanthipeptide class i | 863095 | 956580 | 93485 | Bogorol A | 100% | Brevibacillus laterosporus strain E7593-50 | 91% of genes show similarity |
| **Genome size: 5294 Kbs** | 5 | NRPS-like | 1755504 | 1799178 | 43674 | | | | |
| | 6 | NRPS | 1846107 | 1897176 | 51069 | | | | |
| | 7 | TransAT-PKS | 2146355 | 2212912 | 66557 | Basiliskamide A/B | 81% | Brevibacillus laterosporus DSM 25 | 94% of genes show similarity |
| | 8 | RRE-containing | 2277556 | 2298974 | 21418 | | | | |
| | 9 | Terpene | 2501030 | 2522922 | 21892 | | | | |
| | 10 | NRPS T1PKS | 2528794 | 2670814 | 142020 | Tauramamide | 72% | Brevibacillus laterosporus DSM 25 c | 88% of genes show similarity |
| | 11 | LAP | 2705378 | 2728924 | 23546 | | | | |
| | 12 | NRPS | 2762067 | 2846484 | 84417 | Laterocidine | 63% | Brevibacillus laterosporus strain E7593-50 | 81% of genes show similarity |
| | 13 | NRPS TransAT-PKS | 2891880 | 2989634 | 97754 | Octapeptin C4 | 11% | Brevibacillus laterosporus strain E7593-50 | 77% of genes show similarity |
| | 14 | T3PKS NRPS T1PKS | 3185118 | 3281661 | 96543 | Zwittermycin A | 44% | Brevibacillus laterosporus DSM 25 | 98% of genes show similarity |
| | 15 | Cyclic lactone autoinducer | 3413478 | 3433999 | 20521 | | | | |
| | 16 | Phosphonate | 3654013 | 3667215 | 13202 | | | | |
| | 17 | NRPS NRP mettalophore | 3989058 | 4113653 | 124595 | Ulbactin F/G | 85% | Brevibacillus laterosporus DSM 25 | 81% of genes show similarity |
| | 18 | NRPS | 4220168 | 4267798 | 47630 | Micrococcin P1 | 8% | Brevibacillus sp. 7WMA2 | 100% of genes show similarity |
| | 19 | RiPP like | 4403123 | 4413965 | 10842 | | | | |
| | 20 | NRPS | 4416476 | 4463489 | 47013 | Tyrocidine | 12% | Brevibacillus laterosporus strain 1821L | 8% of genes show similarity |
| **Total genome investment for toxin production** | | | | | 1079559 | | | | |
| | 1 | RRE-containing | 491329 | 511646 | 20317 | Capsular polysaccharide | 13% | Paenibacillus chitinolyticus strain KCCM 41400 | 68% of genes show similarity |
| **Grass 111** | 2 | RiPP like | 602806 | 614992 | 12186 | | | | |
| **Paenibacillus chitinolyticus** | 3 | Crocagin HR-T2PKS | 703032 | 772151 | 69119 | | | | |
| | 4 | Terpene | 2070073 | 2090981 | 20908 | Carotenoid | 33% | Paenibacillus sp. UNC499MF, | 100% of genes show similarity |
| **Genome size: 5844 Kbs** | 5 | TransAT-PKS NRPS | 2276916 | 2353721 | 76805 | Pelgipeptin | 25% | Paenibacillus sp. UNC499MF | 100% of genes show similarity |
| | 6 | TransAT-PKS NRPS T1PKS phosp | 2733765 | 2859075 | 125310 | Zwittermycin A | 22% | Paenibacillus sp. UNC499MF | 98% of genes show similarity |
| | 7 | NRPS NRP-mettalophore | 3020032 | 3141605 | 121573 | Bacillibactin | 100% | Paenibacillus sp. HGH0039 | 82% of genes show similarity |
| | 8 | Opine-like mettlophore | 3149007 | 3171148 | 22141 | Bacillopaline | 100% | Paenibacillus sp. UNC499MF | 90% of genes show similarity |
| | 9 | T3PKS | 3283285 | 3324439 | 41154 | | | | |
| | 10 | Cyclic lactone autoinducer | 4114648 | 4135118 | 20470 | | | | |
| | 11 | Proteusin | 4712421 | 4732660 | 20239 | | | | |
| | 12 | RRE-containing | 5225993 | 5246250 | 20257 | | | | |
| **Total genome investment for toxin production** | | | | | 570479 | | | | |
| | 1.1 | NI siderophore | 323559 | 357072 | 33513 | Schizokinen | 60% | Peribacillus frigoritolerans strain 44 | 92% of genes show similarity |
| **Grass 112** | 1.2 | Thoamitides | 675329 | 697285 | 21956 | | | | |
| **Peribacillus simplex** | 1.3 | Terpene | 714952 | 736847 | 21895 | Citillin A/B | 7% | Peribacillus simplex strain SH-B26 | 88% of genes show similarity |
| | 1.4 | LAP | 1287342 | 1310880 | 23538 | | | | |
| **Genome size: 5249 Kbs** | 1.5 | Terpene | 2161277 | 2182095 | 20818 | | | | |
| | 1.6 | T3PKS | 2329721 | 2370809 | 41088 | Micrococcin P1 | 8% | Peribacillus simplex strain SH-B26 | 92% of genes show similarity |
| | 1.7 | Cyclic lactone autoinducer | 2415104 | 2435858 | 20754 | | | | |
| | 1.8 | NRPS | 2545770 | 2614284 | 68514 | | | | |
| | 1.9 | Guanidinotides | 2704007 | 2726483 | 22476 | | | | |
| | 1.10 | Betalactone | 3150620 | 3234852 | 84232 | Fengycin | 46% | Peribacillus simplex strain SH-B26 | 52% of genes show similarity |
| | 1.11 | NRPS | 3671568 | 3744084 | 72516 | Leupeptin Pr/Ac | 25% | Metabacillus halosaccharovorans strain DSM 253 | 30% of genes show similarity |
| | 1.12 | Ranthipeptide | 4623785 | 4654186 | 30401 | | | | |
| | | | | | 461701 | | | | |
